# DNA-dependent protein kinase promotes DNA end processing by MRN and CtIP

**DOI:** 10.1101/395731

**Authors:** Rajashree A. Deshpande, Logan R. Myler, Michael M. Soniat, Nodar Makharashvili, Linda Lee, Susan P. Lees-Miller, Ilya J. Finkelstein, Tanya T. Paull

## Abstract

The repair of DNA double-strand breaks occurs through non-homologous end joining or homologous recombination in vertebrate cells - a choice that is thought to be decided by a competition between DNA-dependent protein kinase (DNA-PK) and the Mre11/Rad50/Nbs1 (MRN) complex but is not well understood. Using ensemble biochemistry and single-molecule approaches, here we show that the MRN complex is dependent on DNA-PK and phosphorylated CtIP to perform efficient processing and resection of DNA ends in physiological conditions, thus eliminating the competition model. Endonucleolytic removal of DNA-PK-bound DNA ends is also observed at double-strand break sites in human cells. The involvement of DNA-PK in MRN-mediated end processing promotes an efficient and sequential transition from non-homologous end joining to homologous recombination by facilitating DNA-PK removal.

**One Sentence Summary:** DNA-dependent protein kinase, an enzyme critical for non-homologous repair of DNA double-strand breaks, also stimulates end processing for homologous recombination.

## Introduction

DNA-dependent Protein Kinase consists of a catalytic kinase subunit (DNA-PKcs) and the DNA end-binding heterodimer of Ku70 and Ku80 (Ku). Together, these proteins form an end recognition complex (DNA-PK) that binds to DNA double-strand breaks within seconds of break formation (*1*). DNA-PK promotes the ligation of two DNA ends by ligase IV, aided by accessory factors and in some cases accompanied by limited end processing (*1*, *2*). The recognition and repair of DNA breaks by the DNA-PK-associated non-homologous end-joining (NHEJ) machinery is generally considered to be the first and rapid phase of DNA repair, occurring within ~30 minutes of DNA damage, while homologous recombination is specific to the S and G_2_ phases of the cell cycle and occurs over a longer time frame (*3*, *4*). DNA-PK is present at micromolar concentrations in cells (*5*) and binds DNA ends in all cell cycle phases, even during S phase at single-ended breaks (*6*).

The MRN complex regulates the initiation of DNA end processing prior to homologous recombination by catalyzing the initial 5’ strand processing at blocked DNA ends and by recruiting long-range nucleases Exo1 and Dna2 (*4*, *7*). At single-ended breaks during replication, the nuclease activity of Mre11 was shown to be important for removal of Ku (*6*), suggesting that MRN catalytic processing of ends during S phase is important for recombination-mediated repair of single-ended breaks. In addition, the presence of protein blocks on DNA ends has been shown to stimulate MRN(X) endonuclease activity in vitro (*8*–*12*), consistent with the idea that chromatin-bound proteins stimulate MRN activities. Here we considered the possibility that the presence of DNA-PK on DNA ends might regulate MRN-dependent end processing and that this interaction may be a critical component of the choice between NHEJ and homologous recombination pathways in human cells.

## Results

To test for the effect of DNA-PKcs on end processing by the MRN complex, we performed a nuclease assay with purified human MRN, CtIP, Ku, and DNA-PKcs (Fig. S1) on a 197 bp linear double-stranded DNA containing a single radiolabel at the end of one of the 5’ strands. Addition of all the protein complexes together to this substrate resulted in the appearance of a single major cleavage product approximately 45nt in length (Fig. 1A, lane 2). We previously reported MRN-mediated removal of the Ku heterodimer alone, showing that MRN makes an endonucleolytic cleavage at approximately 30 bp from the DNA end (*13*); here we observe that the inclusion of DNA-PKcs in the reaction increases the efficiency of the cutting by approximately 50-fold as well as changing the position of the nuclease cleavage site to a location approximately 45 nt from the end (Fig. 1A, lanes 2 and 3). Importantly, efficient formation of the cleavage product requires CtIP, which is essential for DNA end resection in human cells (*14*). Mre11 nuclease activity in vitro is strictly manganese-dependent (*8*, *15*, *16*); however, Mre11/Rad50 complexes from archaea and from T4 bacteriophage also exhibit nuclease activity in the absence of manganese when physiologically relevant protein cofactors are present (*17*, *18*). Here we observe that human MRN makes endonucleolytic cuts in magnesium-only conditions and that this requires the presence of both Ku and DNA-PKcs (Fig. 1A, lane 10). Since human cells do not contain high levels of manganese (*19*), we conclude that the physiologically relevant endonuclease activity of MRN is thus dependent on DNA-PK.

**Figure 1:**
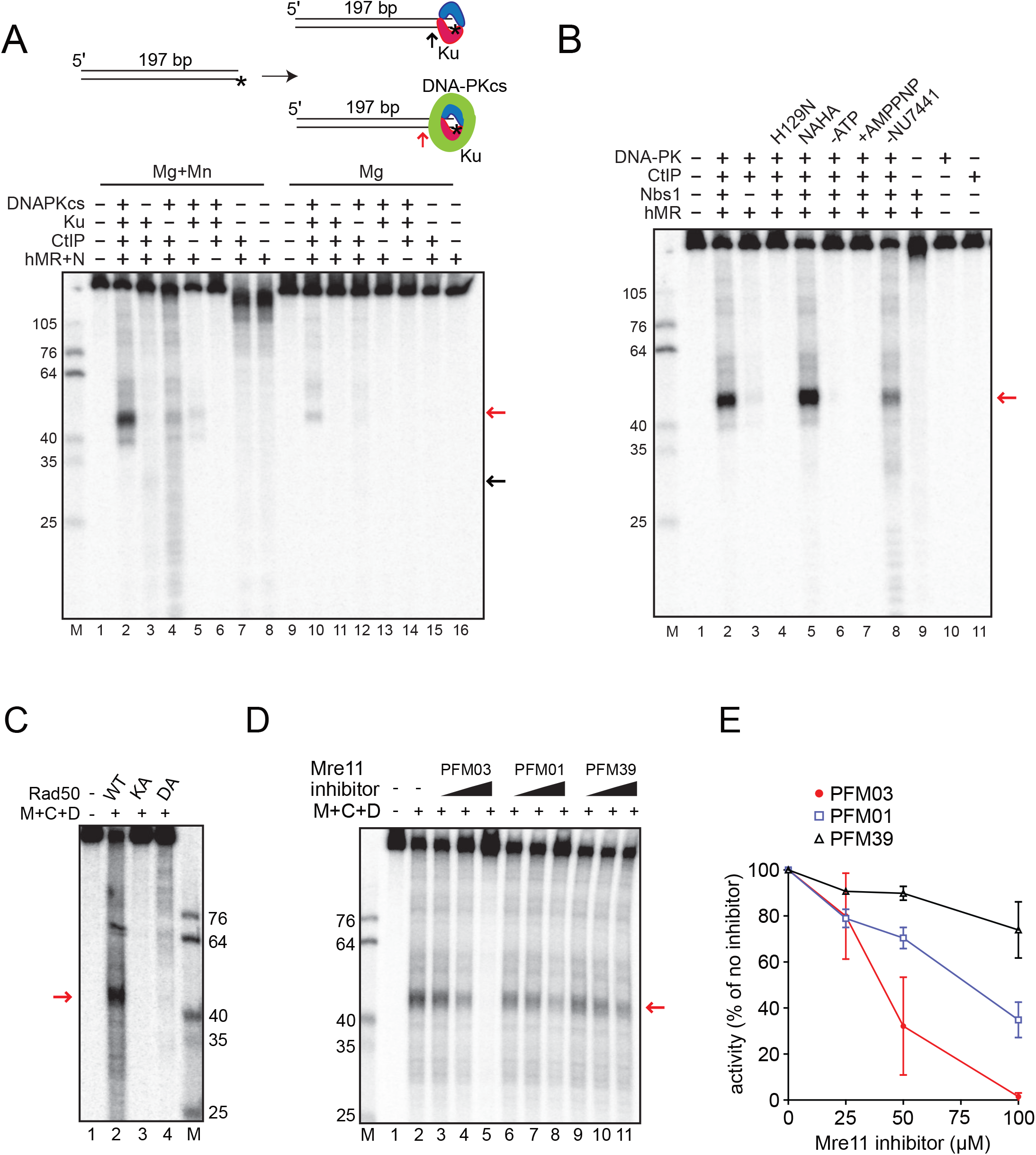
Nucleolytic removal of DNA-PK by MRN and its stimulation by CtIP. (A) Nuclease reactions were performed with a 197 bp DNA substrate, 5’ labeled with ^32^[P](asterisk), with MRN (50 nM), CtIP (80 nM), Ku (10 nM), and DNA-PKcs (10 nM) as indicated, in the presence of both magnesium and manganese (lanes 1-8) or magnesium only (lanes 9-16). All reactions contained the DNA-PKcs inhibitor NU7441. Products were visualized by denaturing polyacrylamide gel electrophoresis and visualized by phosphorimager. Red and black arrows indicate the predominant product in the presence of DNA-PKcs and Ku, or Ku alone, respectively. (B) Nuclease assays were performed as in (A) in the presence of both magnesium and manganese with wild-type MRN or MRN containing nuclease-deficient Mre11 (H129N). Reactions without ATP or with AMP-PNP instead of ATP are indicated, as is the reaction without NU7441. (C) Nuclease assays were performed as in (A) in the presence of magnesium, manganese, ATP, NU7441, CtIP (C), DNA-PK (D) with MRN (M) containing wild-type Rad50 (WT) or ATPase-deficient Rad50 K42A (KA) or D1231A (DA). (D) Nuclease assays were performed as in (A) in the presence of both magnesium and manganese with wild-type MRN in the presence of 25, 50 and 100 μM Mre11 inhibitors and NU7441. (E) Quantitation of the MRN endonuclease observed in presence of Mre11 inhibitors expressed as % of the activity in the absence of inhibitors. Error bars represent standard deviation from two replicates.

Mre11 and CtIP both exhibit endonucleolytic activity (*8*, *20*) so we asked which protein is responsible for the catalysis. We found that substituting wild-type Mre11 with a nuclease-deficient form of Mre11, H129N (*21*, *22*), completely abolished DNA-PK-dependent cutting by MRN, whereas we observed robust activity with nuclease-deficient N289A/H290A CtIP (Fig. 1B, lanes 4 and 5); thus, we conclude that the processing is done by the catalytic activity of Mre11.

We previously observed that ATP binding by Rad50 promotes Nbs1-dependent endonucleolytic cutting by Mre11 (*8*, *15*). Exclusion of ATP or substitution of the nonhydrolyzable AMP-PNP analog for ATP blocks MRN nuclease activity in the presence of DNA-PK (Fig. 1B, lanes 6 and 7), suggesting that ATP hydrolysis by Rad50 is critical for DNA-PK-promoted MRN nuclease activity. We also tested MRN complexes containing Rad50 K42A and D1231A proteins with mutations in the Walker A and Walker B ATP-binding motifs, respectively, that are deficient in ATP hydrolysis (*23*). Both complexes failed to support MRN endonuclease activity (Fig. 1C), confirming the requirement for Rad50 catalytic activity.

Phosphorylation of Ku has been shown to result in the removal of Ku from DNA (*24*), and autophosphorylation of DNA-PKcs also removes DNA-PKcs from the Ku-DNA complex (*25*–*29*), therefore in reactions shown in Fig. 1A and B, we have also included an inhibitor of DNA-PKcs, NU7441, to block its kinase activity. Removal of the inhibitor reduced the efficiency of the reaction (Fig. 1B, lane 8), suggesting that reduced stability of DNA-PK on DNA reduces the nuclease activity of MRN on the DNA-PK-bound substrate. However, the endonucleolytic product is still observed in the absence of the inhibitor (also see Fig. S2), thus blocking DNA-PKcs phosphorylation is not absolutely essential.

Small molecule inhibitors have been developed that specifically block either the exonuclease (PFM39) or the endonuclease (PFM01 and PFM03) activity of Mre11 (*30*). We found that PFM01 and PFM03 reduced MRN endonuclease activity by 60% and 98%, respectively, compared to 25% inhibition by PFM39 (Fig. 1D,E). Efficient inhibition by PFM03 and PFM01 are consistent with Mre11 endonucleolytic activity acting on DNA ends bound by DNA-PK in this assay.

CtIP is phosphorylated by several different kinases in human cells (*31*)(Fig. 2A). Some of these phosphorylation events have been shown to promote DNA end resection (*8*, *9*, *32*–*37*), most notably the CDK-dependent phosphorylation of T847. In addition, ATM/ATR phosphorylation of T859 also plays an important role in CtIP binding to chromatin and CtIP-mediated resection (*38*, *39*). To test the function of CtIP phosphorylation in the MRN endonuclease assay, we purified recombinant CtIP proteins containing phospho-blocking or phospho-mimic mutations at the T847 and T859 sites (Fig. S1). Addition of these mutant proteins to the MRN endonuclease assay with DNA-PK shows that T847 and T859 phosphorylation are critical, as the phospho-deficient T847A and T859A mutants resulted in loss of CtIP stimulation (15- and 3-fold stimulation of MRN, respectively, compared to 43-fold by WT) (Fig. 2B, lanes 5-10, Fig. 2C). We know that there is substantial constitutive phosphorylation of CtIP expressed and purified from insect cells (*40*) and infer from this result that T847/T859 phosphorylated species in the recombinant protein preparation are essential for stimulation of the MRN endonuclease activity. In contrast to these sites, phosphorylation at S327 or the ATM sites S231, S664 and S745 appears not to be important for the endonucleolytic activity (Fig. 2B, lanes 11-13). We previously reported that the ATM phosphorylation sites are important for CtIP intrinsic nuclease activity (*40*). Here we show that these sites do not regulate DNA-PK-dependent Mre11 endonuclease activity, consistent with our observation that the N289A/H290A mutations that reduce CtIP nuclease activity also do not impact Mre11 function in this assay.

**Figure 2:**
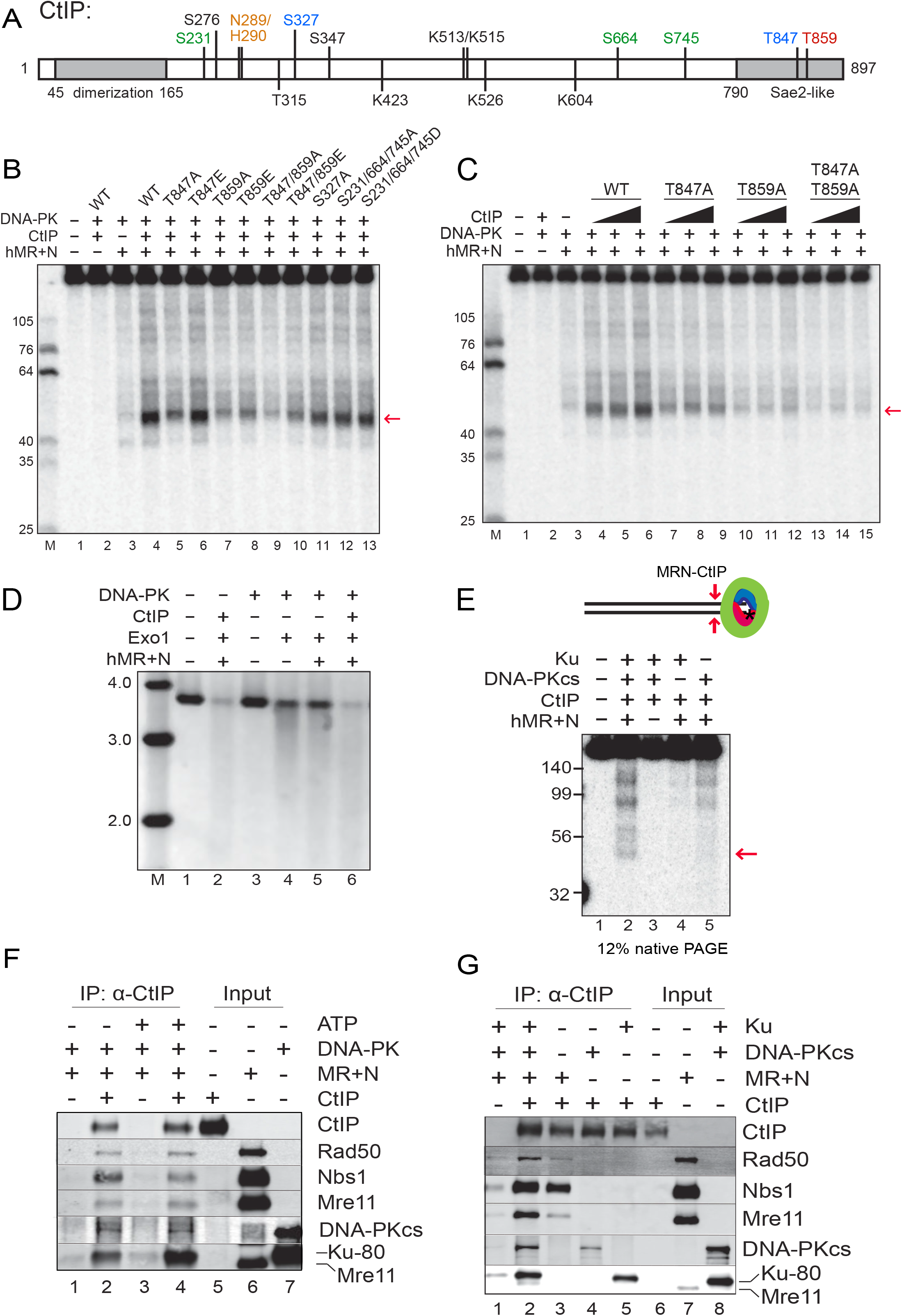
MRN, DNA-PK, and CtIP promote double-stranded DNA end resection. (A) A linear map of CtIP indicating a subset of known phosphorylated residues as well as residues important for DNA binding and catalytic activity as discussed in the text (orange: sites required for nuclease activity, green: ATM-dependent phosphorylation sites, blue: CDK-dependent phosphorylation sites, red: ATR/ATM-dependent phosphorylation site). (B) Nuclease assays were performed with MRN (12.5 nM), CtIP (40 nM), Ku (10 nM) and DNA-PKcs (10 nM), and NU7441 as in Fig. 1A but with various mutants of CtIP as indicated, in the presence of both magnesium and manganese. The red arrow indicates the predominant product formed in the presence of DNA-PKcs. (C) Nuclease assays were performed as in (B) with titrations of CtIP (10, 20 and 40 nM). (D) DNA end resection on a plasmid substrate (3.6 kb) was performed with MRN, CtIP, DNA-PK, and Exo1 as indicated, in the presence of DNA-PKcs inhibitor. Reaction products were separated in a native agarose gel, which was stained with SYBR green; MW ladder migration is shown (kb). (E) Double-stranded DNA cleavage products from the MRN nuclease assay with DNA-PK and CtIP were detected on a 12% native polyacrylamide gel. The red arrow indicates the ~45 bp product. (F) Protein-protein interactions between CtIP, MRN, and DNA-PK were measured by immunoprecipitation with anti-CtIP antibody in the presence or absence of ATP followed by western blotting of bound proteins, as indicated. (G) Interactions between CtIP, MRN, DNA-PKcs and Ku was measured with CtIP immunoprecipitation as in (F) in the presence of ATP.

While mutation of the CtIP T847 residue to phospho-blocking alanine inhibited its stimulation of Mre11 nuclease activity, the phospho-mimic T847E mutant completely restored this ability in vitro (Fig. 2B, lane 6). This is important as the T847E allele of CtIP was previously shown to restore localization of CtIP to sites of DNA damage as well as the resection of doublestrand breaks, and promoted efficient CDK-dependent processing of these breaks (*41*). Thus Mre11 endonuclease activity is linked with the cell cycle-dependent processing of DNA damage via the phosphorylation of T847 in CtIP, and will therefore only occur efficiently during the S and G_2_ phases of the cell cycle. The T859E phospho-mimic version of CtIP only partially restores Mre11 activity (Fig. 2B, lane 8), also consistent with the initial study of T859 phosphorylation by ATR in Xenopus extracts which reported that the T859E (T818E in Xenopus CtIP) is only weakly active in supporting double-strand break resection in CtIP-depleted extracts (*38*). Overall, however, the combined data suggest that CDK and ATR/ATM-mediated phosphorylation of CtIP on T847 and T859 are critical for stimulation of MRN endonuclease activity on DNA-PK-bound ends.

Exo1 is one of the critical long-range exonucleases that performs DNA end resection in human cells (*4*) and we have previously shown a positive effect of MRN on Exo1-mediated degradation of DNA (*13*, *42*, *43*). Despite the well-known importance of CtIP in resection in human cells, however, we have not observed any effects of recombinant CtIP on Exo1 activity in previous experiments. Here we examined the effect of CtIP on Exo1 on a plasmid DNA substrate in the presence of MRN and DNA-PK (Fig. 2D). As expected, Exo1 is substantially blocked by the presence of DNA-PK in the reaction, an inhibition that is relieved by the inclusion of MRN and requires CtIP (Fig. 2D, lanes 4-6). This finding that CtIP stimulates the activity of Exo1 7-fold in the presence of DNA-PK is consistent with the effect of CtIP on Mre11 nuclease activity we observe on the labeled substrates (Fig. 1).

We recently showed that MRN can remove DNA end blocks by sequential endo-exo-endo activities, generating a double-strand break adjacent to the block (*8*, *13*), a model first suggested by processing of meiotic DNA breaks (*44*). In the presence of DNA-PK bound to DNA ends, we found that the MRN complex executes similar end-processing of the radiolabeled substrate to generate a new double-strand break. We detected a ~ 45 bp dsDNA product using native PAGE (Fig. 2E, lane 2), corresponding to the major product seen on denaturing gels (Fig. 1). This activity was dependent on MRN and was enhanced when both Ku and DNA-PKcs were present in the reaction (Fig. 2E, lane 2). However, the formation of the dsDNA product was less efficient (approximately 1 to 2% of the substrate cleaved at site proximal to the radiolabel) compared to the single-strand endonucleolytic product seen on denaturing gels (approximately 27 to 34% of the substrate cleaved adjacent to the labeled 5’ end)(Fig. 1A, B).

The unexpected dependence of Mre11 endonuclease activity on DNA-PK suggests that there may be specific protein-protein interactions underlying the cooperative behavior. We tested this by immobilizing purified, human CtIP on magnetic beads and monitoring binding of DNA-PK or MRN. Both MRN and DNA-PK bound to CtIP in an ATP-independent manner (Fig. 2F). Binding analysis of Ku and DNA-PKcs separately showed that each of the components of DNA-PK has affinity for CtIP (Fig. 2G). Interestingly, binding of Nbs1 (added to this reaction separately from the MR complex) to CtIP is mostly independent of DNA-PK while binding of MR to CtIP improves 7 to 8-fold with DNA-PK present. Addition of EtBr or Benzonase to the reaction did not affect the interaction, confirming that the binding is independent of DNA (Fig. S3A). Although the stimulation of Mre11 nuclease activity requires CtIP phosphorylation on T847 and T859, we did not observe any obvious deficiency in binding between CtIP, MRN, and DNA-PK with the T847A/T859A CtIP mutant protein (Fig. S3B), thus the effect of the phosphorylation does not appear to be manifested through CtIP association with the other factors.

### Single-molecule observations show MRN/CtlP-mediated removal of DNA-PK and MRN from DNA ends

To examine the interactions between DNA-PK and the MRN complex in greater detail, we turned to single-molecule microscopy. In this “DNA curtains” assay, arrays of lambda DNA molecules (~48 kb long) are attached via a biotin-streptavidin linkage to a microfluidic flowcell that is also passivated with a fluid lipid bilayer (Fig. 3A) (*45*). Microfabricated barriers align the DNA molecules and organize them for high throughput analysis of bound proteins, which we have previously used to examine the interplay between the MRN complex and Ku at individual DNA ends (*13*). Here we adapted this assay to observe the behavior of the DNA-PK complex, with DNA-PKcs labeled using a DNA-PKcs-specific antibody. Injection of fluorescently labeled DNA-PKcs led to only transient association with the DNA, as indicated by non-specific sliding in the direction of buffer flow and subsequent release from the DNA ends (half-life = 5 ± 3 sec, N=17; Fig. 3B, C). Given that formation of the DNA-PK complex requires Ku and DNA (*46*), we considered that pre-loaded Ku may stabilize DNA-PKcs on the DNA ends. To test this, we injected HA-tagged Ku heterodimer onto the DNA curtains, labeled with anti-HA antibody. As we observed previously, Ku bound only to DNA ends and did not slide on the DNA with buffer flow (*13*). Subsequent injection of DNA-PKcs did result in more stable association of the kinase with the Ku-bound ends; however, the inclusion of ATP in the buffer promoted rapid loss of ~80% of the bound Ku molecules (Fig. 3D-F). These are single-turnover reactions that we observe with constant buffer flow, so even transient release of DNA-PKcs from the DNA results in loss of association. We considered that the ATP-dependent release might be associated with the phosphorylation of Ku70 by DNA-PKcs, which inhibits DNA binding by Ku (*24*). To prevent Ku phosphorylation and release by DNA-PKcs, we purified a phospho-blocking mutant of Ku(T305A/S306A/T307A/S314A/T316A, “5A”), which inhibits DNA-PKcs-induced release of Ku (*24*). Ku(5A) bound to DNA for >2000 seconds, consistent with our previous observations of extremely stable binding of wild-type Ku (Fig. 3E, F)(*13*). Ku(5A) stabilized DNA-PKcs at the DNA ends even in the presence of ATP (Fig. 3D-F), confirming that phosphorylation of the Ku70 S/TQ 305-316 cluster controls the association of DNA-PK with DNA ends.

**Figure 3:**
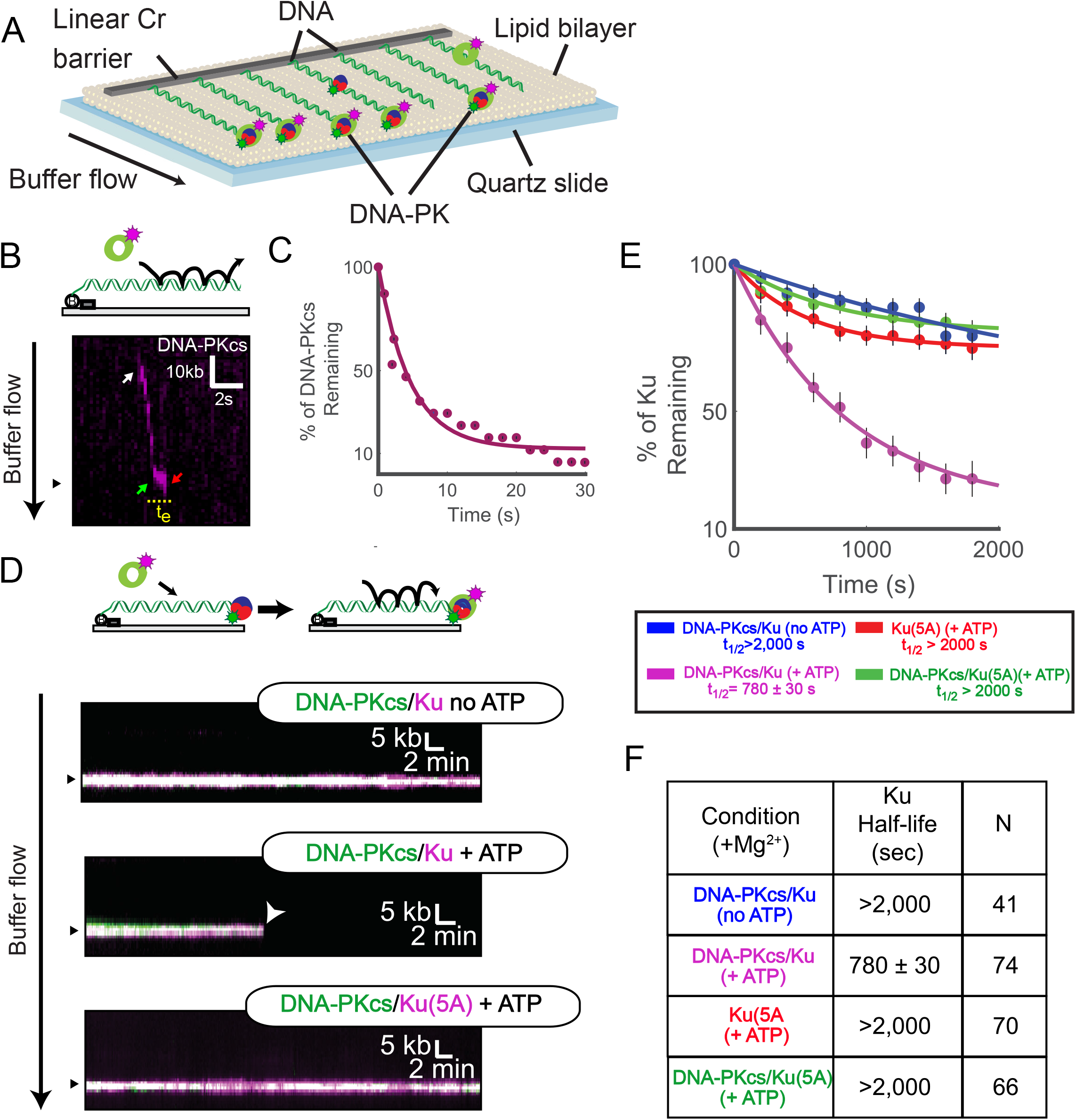
Single-molecule visualization of DNA-PK on DNA. (A) Schematic of the DNA curtains assay for DNA-PK. (B) Illustration and kymograph (time series of one molecule over the course of the reaction) of DNA-PKcs injection onto DNA curtains. White arrow indicates a single DNA-PKcs binding event. This molecule then slides along the DNA in the direction of buffer flow to reach the DNA end (green arrow). The molecule then associates for a short time at the end (t_e_) before dissociation (red arrow). (C) Lifetime of DNA-PKcs on DNA curtains in the presence of ATP. (D) Illustration and kymographs of DNA-PKcs co-localizing with an end-bound Ku and Ku(5A) in the presence or absence of ATP as indicated. (E and F) Lifetime of Ku (WT or 5A) on DNA curtains in the presence or absence of DNA-PKcs or ATP as indicated. Table shows half-life of Ku(WT) or Ku(5A) under various conditions and the number of molecules observed (N).

We next observed the interplay between MRN and DNA-PK (DNA-PKcs with Ku(5A)) using the DNA curtains platform. Consistent with our ensemble biochemical assays (Fig. S2), injection of dark (unlabeled) MRN and CtIP onto DNA curtains with both fluorescent DNA-PKcs and fluorescent Ku in the presence of a physiologically relevant cleavage buffer (5 mM Mg^2+^) led to removal of both DNA-PKcs and Ku from the ends (Fig. 4A).

**Figure 4.**
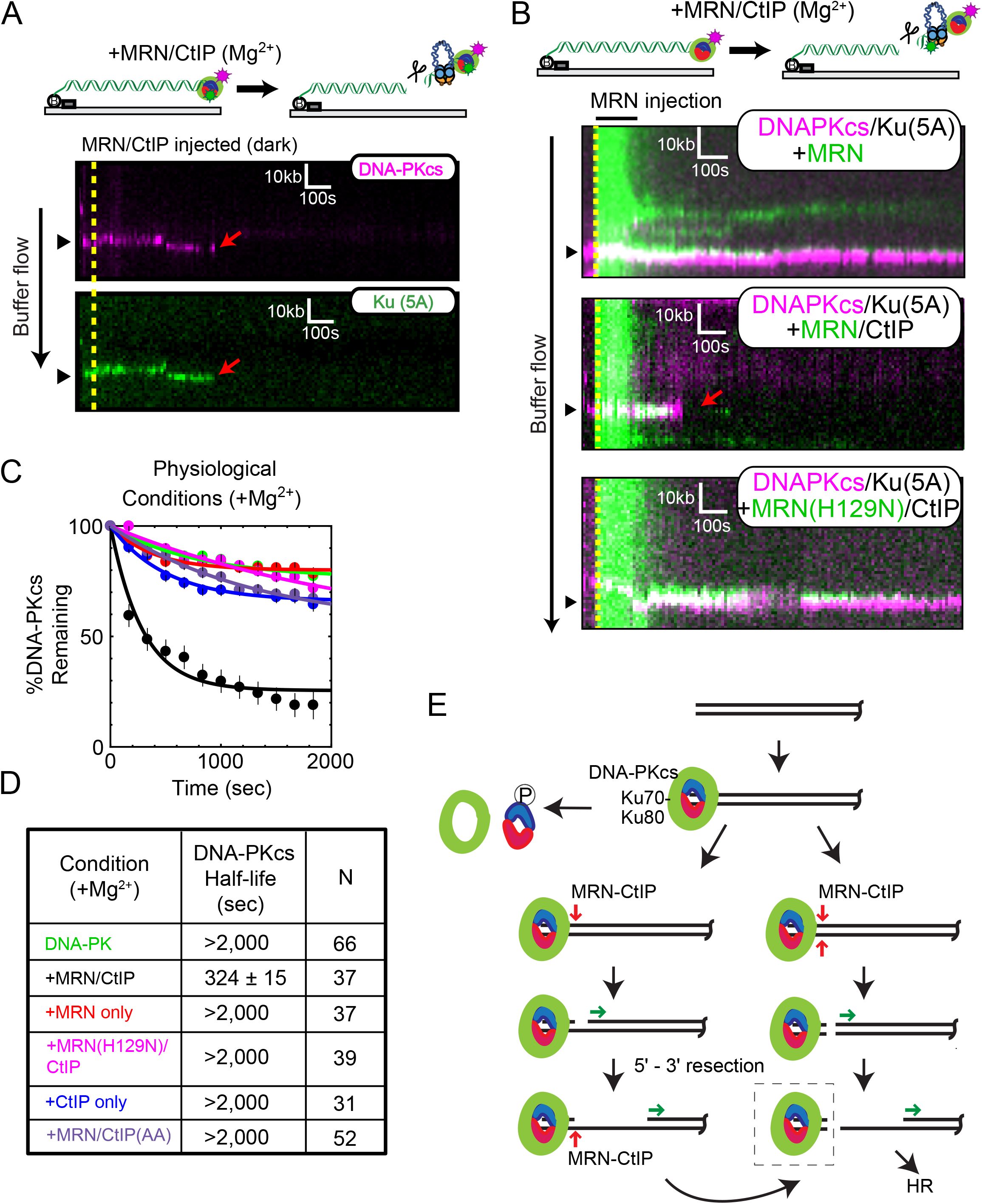
Single-molecule visualization of DNA-PK removal by MRN and CtIP. (A) Illustration and kymographs of DNA-PKcs (magenta) and Ku (green) upon injection of MRN and CtIP (both unlabeled (dark)). (B) Kymographs of DNA-PKcs (magenta) with Ku (unlabeled), with injection of MRN (green) alone (top), or MRN with CtIP (middle), or a nuclease-dead mutant of MRN (H129N) with CtIP (bottom). (C and D) Lifetimes and associated half-lives of the DNA-PK complex (as observed by DNA-PKcs occupancy) upon no injection (green), injection with MRN and CtIP (black), MRN alone (red), nuclease deficient H129N MRN and CtIP (magenta), CtIP only (blue), or MRN with phospho-blocking T847A/T859A CtIP (purple). Table shows half-life of DNA-PKcs under various conditions and the number of molecules observed (N). (E) Schematic model for DNA-PK removal from double-strand break ends, as discussed in the text. DNA-PKcs binds to the Ku heterodimer bound to DNA at the DSB. Transition from NHEJ to HR results from 1) DNA-PKcs phosphorylation of Ku, resulting in dissociation of Ku as well as DNA-PKcs from ends, 2) DNA-bound DNA-PK stimulation of MRN single-strand endonucleolytic cleavage followed by 5’ - 3’ resection, or 3) DNA-PK stimulation of MRN double-stranded endonucleolytic cleavage resulting in DNA-PK loss and 5’ - 3’ resection. Long range 5’ - 3’ resection ensues from the nick or a new double-strand break, creating 3’ single-stranded DNA that is used for homology search. The DNA-bound DNA-PK complex (dashed box) is the species isolated in the modified ChIP protocol (Fig. 5 and Fig. S4).

In order to probe the dynamics of this activity, we utilized fluorescently-labeled MRN complex together with fluorescent DNA-PKcs (Ku is unlabeled in this case) and measured the occupancy of DNA-PK on ends that are also bound by MRN. Injection of MRN alone led to stable co-localization of the two complexes (DNA-PK half-life >2000 sec), but injection of MRN and CtIP together led to removal of both Ku and DNA-PKcs from the DNA ends (half-life = 324 ± 15 sec, N=37; Fig. 4B-D). In contrast, an MRN nuclease-deficient mutant (H129N) with CtIP did not remove DNA-PK; neither did CtIP added in the absence of MRN. Furthermore, injection of MRN with CtIP containing phospho-blocking mutations T847A and T859A also failed to remove DNA-PK (Fig. 4B-D). This data suggests that co-localization of MRN with the DNA-PK complex is not sufficient to facilitate removal and that, consistent with our ensemble assays, phosphorylated CtIP is required for DNA-PK removal by MRN nuclease activity. It is notable that the rate of DNA-PK removal by MRN/CtIP under these conditions (t_1/2_ = 324 ± 15 sec) is faster than the removal of Ku by DNA-PK-mediated phosphorylation (t_1/2_ = 780 ± 30 sec), thus the MRN-mediated pathway is likely to occur in a time-frame comparable to other pathways of Ku removal in cells.

Our previous studies showed that MRN could remove Ku from DNA ends via a manganese-dependent reaction (*8*, *13*). Here we observe approximately the same kinetics of DNA-PK release catalyzed by MRN and CtIP as we did previously for Ku, but in magnesium-only conditions (all of the single-molecule experiments shown here were performed in the absence of manganese). These results demonstrate that DNA-PK stimulates Mre11 nuclease activity in the absence of manganese in physiologically relevant conditions and that CtIP is an essential component of this reaction.

These observations suggest a conserved mechanism for removal of blocks from DNA ends by the MRN complex (model in Fig. 4E). At two-ended double-strand breaks, NHEJ would first engage the ends, and in cases where the ends are compatible, the ends would undergo productive end-joining. In situations where NHEJ fails and the DNA-PK complex remains at the DSB ends, Ku could be removed by DNA-PK-mediated phosphorylation (*24*). Alternatively, MRN can remove the complex through endonucleolytic processing, a reaction which is greatly enhanced by phosphorylated CtIP and is dependent on DNA-PK, as we show in this study. MRN activity nicks either one or both the strands close to the DNA-PK complex in a sequential or simultaneous manner. The nicked or freed end is then used for extensive 5’ to 3’ resection by exonucleases allowing damage repair through homologous recombination or homology-dependent mechanisms.

### MRN nuclease-mediated cutting of DNA-PK-bound ends occurs in human cells

To determine if nucleolytic removal of DNA-PK occurs in human cells, we performed a chromatin immunoprecipitation (ChIP) assay using the inducible AsiSI restriction enzyme system in U2OS osteosarcoma cells (*47*). We modified the standard ChIP protocol (Fig. S4) by reducing the duration of formaldehyde crosslinking and eliminating the extensive sonication used to fragment DNA. This gentle lysis procedure, which we are naming “Gentle Lysis And Size Selection-ChIP”, or GLASS-ChIP, allowed us to specifically separate small (50 to 300 bp) DNA-PK-associated DNA fragments from the bulk chromatin. The fragments were then further purified and used for quantitative PCR analysis. We monitored four genomic AsiSI sites (*48*) and found that this procedure generated DNA-PK-associated DNA fragments at all four sites that were strongly dependent on 4-OHT addition (Fig. 5A).

**Figure 5.**
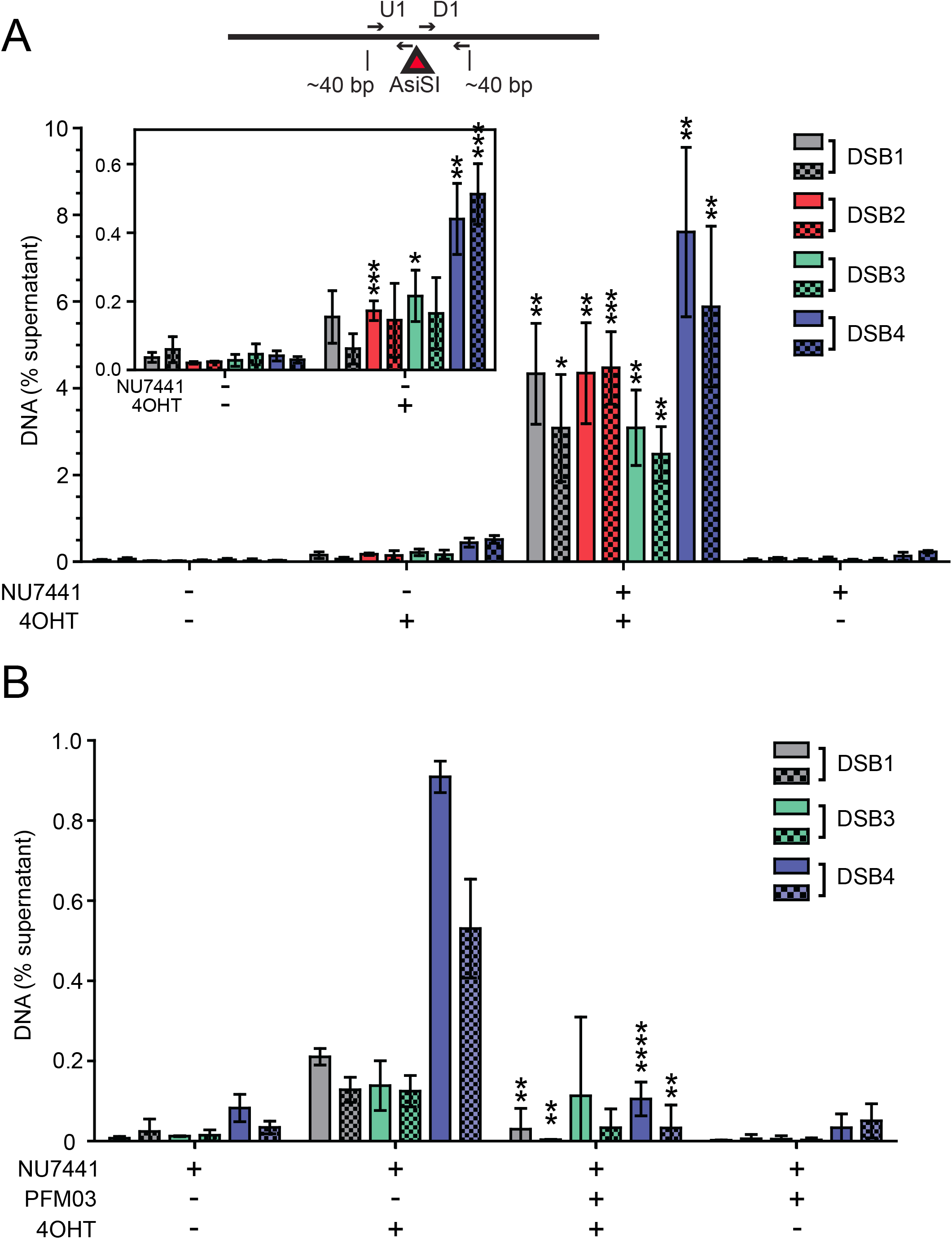
DNA-PK-associated DNA end fragments are observed at AsiSI breaks in human cells using GLASS-ChIP. (A) Small dsDNA products resulting from nucleolytic cleavage of DNA-PK bound AsiSI-generated DNA ends (dashed line box in A) were isolated from U2OS cells treated with 4-OHT or vehicle for 4 h as indicated. NU7441 (10 μM) was added as indicated for 5 h starting at 1 h before 4-OHT addition. DNA-PK-bound DNA was isolated using a modified ChIP protocol (GLASS-ChIP, Fig. S4) and quantified by qPCR using primers located ~30 nt from the AsiSI cut site (results from primers ~300 nt from cut sites in Fig. S5). Primer set U1 (solid) is upstream whereas D1 (checkered) is downstream of the AsiSI cut sites. The DNA quantitated from U2OS cells in the presence or absence of 4-OHT to induce AsiSI and DNA-PKcs inhibitor (NU7441) is shown for each AsiSI site (*48*). The inset magnifies the results for experiments performed in the absence of NU7441. Results shown are from 3 independent biological replicates, with student 2-tailed T test performed; *, **, and *** indicate p < 0.05, 0.01, and 0.001, respectively, in comparison to equivalent samples without 4-OHT. (B) The GLASS-ChIP protocol was performed as in (A) using cells treated with DNA-PKcs inhibitor (NU7441, 10 μM), Mre11 inhibitor (PFM03, 100 μM), and 4-OHT for 1 h as indicated. Results shown are from 3 independent biological replicates, with student 2-tailed T test performed; ** and **** indicate p < 0.005 and 0.0001, respectively, in comparison to equivalent samples without PFM03.

When cells were exposed to DNA-PKcs inhibitor NU7441 during AsiSI induction, quantitation of DNA located very close (~30 nt) to the AsiSI genomic sites showed a 25-250-fold increase over background levels (i.e. levels of product formed with NU7441 but in the absence of 4-OHT induction)(Fig. 5A), consistent with the nucleolytic removal of DNA-PK-bound DNA ends we observed in vitro. These levels dropped significantly when measuring sites located farther away (~300 nt) from the AsiSI cut site (Fig. S5), and no signals above background were observed at representative locations distant from AsiSI sites. With DNA-PKcs inhibitor present as it is here, we expect that NHEJ is blocked and MRN cleavage of DNA-PK-bound ends is maximal, as we observed in purified protein reactions (Fig. 1 and 2).

Induction of AsiSI with 4-OHT in the absence of DNA-PKcs inhibitor also generated DNA in the GLASS-ChIP assay, approximately 3 to 160-fold higher than background depending on the site, measured with primers 30 nt from the AsiSI location (Fig. 5A inset). Under these conditions NHEJ is not blocked, thus the release of DNA-PKcs with associated DNA is expected to occur only as a consequence of DSB processing. Importantly, the observation of these products in the absence of DNA-PKcs inhibitor shows that processing of DNA-PKcs-bound ends occurs in human cells under normal physiological conditions.

To determine if the DNA-PK-bound products arise through Mre11 nuclease activity we treated the cells with endonuclease inhibitor PFM03 based on our in vitro observations (Fig. 1D-E). In preliminary experiments, we found that addition of 4-OHT to induce AsiSI activity in cells also exposed to PFM03 (100 μM) and DNA-PKcs inhibitor NU7441 resulted in complete cell death within 1.5 h. To circumvent this, we limited the treatment with 4-OHT and both DNA-PKcs and Mre11 inhibitors to 1 hour. In the absence of the Mre11 inhibitor, we observed short DNA fragments at 3 of 4 of the genomic loci tested that were dependent on 4-OHT (Fig. 5B), albeit at 10- to 20-fold reduced levels compared to the 4 hour 4-OHT induction (Fig. 5A). With the addition of PFM03, there was a significant reduction in the recovery of these fragments, indicating that Mre11 nuclease activity is indeed responsible for the creation of DNA-PK-bound DSB products (Fig. 5B).

## Discussion

Here we have shown that DNA-PK plays an important role in DNA end processing through its stimulation of Mre11 endonuclease activity. The extremely high concentration of DNA-PK in mammalian cells, particularly in human cells (*5*), and the high affinity of Ku for DNA ends (*49*) means that double-strand break ends will be immediately bound by DNA-PK (see model in Fig. 4E). Removal of this block can be promoted by several distinct mechanisms: by DNA-PKcs phosphorylation of Ku70 which reduces its DNA-binding affinity (*24*), by ubiquitination of Ku and degradation by proteasome-dependent or independent mechanisms (*50*–*52*), or single-strand or double-strand endonucleolytic incision of DNA by MRN as we show in this study. Single-strand DNA cutting by Mre11 followed by 3’ to 5’ Mre11 exonuclease activity was demonstrated as a mechanism for Spo11 removal by Mre11-Rad50-Xrs2 (MRX) in yeast (*44*).

We also demonstrate in this work that double-strand DNA cutting by MRN is promoted by stable occupancy of DNA-PK on the ends, as shown by the increased efficiency of MRN cleavage in the presence of the DNA-PK inhibitor in vitro and by our observation of the release of DNA-PK-bound DNA fragments in human cells. This requirement for stable occupancy of DNA-PK ensures that MRN does not cleave ends which are undergoing successful end-joining, as this outcome also leads to DNA-PKcs dissociation (*1*). DNA-PK and MRN have often been viewed as competitors for DNA ends, but the data presented here shows that these complexes are linked in an ordered pathway, consistent with cytological observations (*53*). The dual roles of DNA-PKcs ensure rapid repair by NHEJ as well as an efficient transition to homologous recombination by promoting MRN-dependent initiation of end processing.

Loss of DNA-PKcs in mice results in radiosensitivity as well as immunodeficiency due to a failure to complete V(D)J recombination, and naturally-occurring mutations in the gene have generated viable but immunocompromised animals (*54*). In contrast, no human patients have been identified with complete loss of DNA-PKcs, and human cell lines lacking the protein exhibit a severe growth deficit in addition to DNA damage sensitivity, extreme genomic instability, and telomere shortening (*55*). Similarly, Ku70 and Ku80-deficient mice are radiosensitive but viable, yet deletion of Ku subunits in human cells generates spontaneous DNA breaks, telomere defects, and cell lethality within several divisions (*56*). This combined set of observations shows that DNA-PK plays many roles in human cells and is more essential for the maintenance of genomic stability in human cells compared to rodents or other mammals. The function of DNA-PKcs in regulating MRN/CtIP-mediated end processing that we describe in this work is likely an important component of the responsibilities of DNA-PKcs in genome maintenance. Further structural analysis is clearly necessary to fully understand how phosphorylated CtIP enables the MRN complex to breach the otherwise impenetrable block that DNA-PK creates on DNA ends and how the many post-translational modifications on CtIP, MRN, and other factors present at ends can modulate these outcomes. The extreme sensitivity of U2OS cancer cells to simultaneous use of DNA-PKcs and Mre11 inhibitors points towards a potential use of these small molecule inhibitors in treatment of cancers with active homologous recombination pathways.

## Materials and Methods

### Plasmid, strains and Protein purification

Recombinant human proteins (MRN wild-type, M(H129N)RN, MR, MR(K42A), MR(D1231A), Nbs1, CtIP wild-type, and CtIP mutants) were expressed and purified from Sf21 insect cells as described previously (*8*, *23*, *40*). 3XFLAG-MRN, 3XHA-Ku70/80 heterodimer, and Exo1-biotin were made as described previously (*13*). DNA-PKcs from human cells was purified as described previously (*57*). The Ku70(5A) mutant (T305A/S306A/T307A/S314A/T316A) was made from the baculovirus transfer vector containing wild-type Ku70 by QuikChange mutagenesis (Stratagene) to create the 5A mutant expression plasmid pTP4143 and the corresponding bacmid pTP4177, which was used with wild-type Ku80 baculovirus to infect Sf21 cells and purify Ku(5A).

### DNA substrates

TP542 (GGTTTTCCCAGTCACGACGTTG) was 5’ [^32^P] radiolabeled using T4 polynucleotide kinase (NEB) and purified (NEB Nucleotide Removal Kit). The 197 bp DNA substrate labeled on the 5’ end was created by PCR using 5’ [^32^P] radiolabeled TP542 (GGTTTTCCCAGTCACGACGTTG) and TP4373(TGGGTCAACGTGGGCAAAGATGTCCTAGCAATGTAATCGTCTATGACGTT) with pTP3718 followed by purification on a native 0.7% agarose gel as described previously (*8*). The DNA substrate used in Figure 2A was made by digesting pCDF-1b plasmid with SspI to create a 3.6 kb linear DNA with blunt ends, and was used without further purification. The lambda DNA substrate used in single molecule studies was prepared as described previously (*13*).

### Nuclease assays

10 μl nuclease assays contained Ku70/80 (10 nM), DNA-PKcs (10 nM), hMR (50 nM), Nbs1 (50 nM) and CtIP (80 nM) unless indicated otherwise in figure legends. Reactions were performed with 0.1 nM DNA in 25 mM MOPS pH 7.0, 20 mM Tris pH 8.0, 80 mM NaCl, 8 % glycerol, 1 mM DTT, 1 mM ATP, 0.2 mM NU7441, 5 mM MgCl_2_, 1 mM MnCl_2_ and 0.2 mg/ml BSA, with additions or exceptions as noted in the figures and legends. hMR and Nbs1 were pre-incubated (10 min at 4 °C) prior to addition to reactions, and Ku and DNA-PKcs were allowed to bind DNA in the presence of NU7441 for 5 min. at 25 °C followed by the addition of other components. Assays were carried out in Protein Lo-Bind eppendorf tubes (Millipore) at 37°C for 30 min. Reactions were stopped with 0.2 % SDS and 10 mM EDTA, lyophilized, dissolved in formamide, boiled at 100°C for 4 min, loaded on denaturing polyacrylamide gels containing 16% acrylamide, 20% formamide and 6M urea, and separated at 40W for 1.5 h, followed by phosphorimager analysis. Mre11 inhibitors were added at 25, 50 and 100 μM final concentration in assay reactions. Mre11 inhibitor dilutions were made in 20% DMSO in Buffer A, which results in 6% DMSO in these assay reactions, and an equivalent concentration of DMSO was used in control reactions. For the detection of dsDNA products, reactions were treated with 10 μg Proteinase K at 37°C for 1h after stopping the nuclease reaction. The samples were then separated using 12% native PAGE in 1X Tris-Borate-EDTA (TBE) buffer at 150V followed by phosphorimager analysis.

DNA resection assays on plasmid substrates were carried out similarly using 0.1 nM 3.6 kb pCDF-1b plasmid linearized with SspI and 1 nM Exo1 in the presence of the DNA-PKcs inhibitor NU7026 (200 μM). Reactions were stopped with 0.2 % SDS and 10 mM EDTA, 1.5 μg Proteinase K was added and incubated at 37°C for 90 min followed by 10 min incubation at 50°C. These reactions were separated in a 0.7% native agarose gel in 1X TBE buffer, stained with SYBR green and visualized with a Chemidoc Gel Imager (Bio-Rad).

### Protein- protein interactions

Immunoprecipitation assays were performed with anti-CtIP antibody (Active Motif 14E1) in Protein Lo-Bind eppendorf tubes (Millipore). In a 40 μl reaction volume, 20 nM Ku70/80, 20 nM DNA-PKcs, 25 nM hMR, 25 nM Nbs1 and 16 nM CtIP were incubated as indicated in the absence of DNA in 25 mM MOPS pH 7.0, 20 mM Tris, pH 8.0, 80 mM NaCl, 8 % glycerol, 1 mM DTT, 1 mM ATP, 0.2 mM NU7441, 5 mM MgCl_2_ and 0.2 mg/ml BSA at 37°C for 10 min. The reaction was then incubated with 2 μg anti-CtIP antibody, 0.4% 3-[(3-Cholamidopropyl)dimethylammonio]-1-propanesulfonate (CHAPS) and 2 μl Protein A/G magnetic beads (Pierce) at 37°C for 30 min with shaking. Beads were isolated using a magnet and washed 3 times with wash buffer (Buffer A (25 mM Tris pH 8.0, 100 mM NaCl, 10% glycerol) containing 0.4% CHAPS and 0.1 mg/ml BSA). Tubes were changed at the third wash step and proteins bound to the beads were eluted with 1X SDS-PAGE loading dye prepared in wash buffer. Samples were heated at 100°C for 5 min and separated by 8% SDS-PAGE followed by Western blotting. Since Mre11 and Ku80 migrate very close to each other, blots were developed sequentially, first for FLAG tagged CtIP, Nbs1 and Mre11, then for DNA-PKcs and His tagged Rad50 and Ku70, and lastly for biotin-tagged Ku80. Proteins were detected using rabbit anti-FLAG (Cell Signaling), rabbit anti-DNA-PKcs (Bethyl Laboratories), and mouse anti-His (Genscript) antibodies as well as streptavidin-Alexa Fluor680 conjugate (Invitrogen) using an Odyssey Imager (Li-Cor).

### Single molecule experiments

Single-molecule DNA curtains were performed as previously described (*13*). For labeling of DNA-PKcs, 400 nM anti-mouse secondary QDots (Fisher) were incubated with 360 nM anti-DNA-PKcs antibody (Abcam ab44815) on ice for 10 minutes in 5 μl DNA-PKcs loading buffer (40 mM Tris-HCl pH 8.0, 0.2 mg/ml BSA, 2 mM MgCl_2_ 2 mM DTT). DNA-PKcs was added to this mixture to a final concentration of 267 nM QDots, 240 nM antibody, and 167 nM DNA-PKcs in 7.5 μl for another 10 minutes on ice. This mixture was diluted to a final volume of 200 μl (10 nM QDots, 9 nM antibody, 6.25 nM DNA-PKcs). Ku(5A) was loaded on DNA curtains essentially as described previously for wild-type Ku (*13*). Next, DNA-PKcs was injected onto DNA curtains from a 100 μl loop at 200 μl/min in DNA-PKcs loading buffer. For MRN cleavage experiments, after DNA-PKcs loading the buffer was switched to MRN cleavage buffer (40 mM Tris-HCl pH 8.0, 60 mM NaCl, 0.2 mg/mlBSA, 5 mM MgCl_2_, 1 mM ATP, 2 mM DTT). After buffer switch, mixtures of 3 nM MRN, 14 nM CtIP, or both were loaded as previously described (*13*).

### GLASS-ChIP assay

Human U2OS cells with inducible AsiSI (*47*) were grown to 50-60% confluency in 150 mm dishes and treated with 600 nM 4OHT for 4 h at 37°C. When used, 10 μM NU7441 was added 1h prior to 4OHT addition. After 4OHT treatment, cells were fixed with 1% formaldehyde for 7 min at RT with gentle rotation. Crosslinking was stopped by addition of 125 mM glycine for 5 min, washed twice with cold PBS, harvested, flash frozen in liquid nitrogen and stored at −80°C. For the ChIP assay, we modified a standard protocol (Abcam) with a gentle lysis procedure and minimal, low-level sonication to rupture cells without extensive DNA damage. The formaldehyde fixed cells were thawed at RT for 5 min, resuspended in RIPA buffer (50 mM Tris-HCl pH8.0, 150 mM NaCl, 2 mM EDTA pH8.0, 1% NP-40, 0.5% Sodium Deoxycholate, 0.1% SDS) with 1x protease inhibitors (Pierce #A32955) and sonicated using a Cell Ruptor at low power, for 10 sec followed by 10 pulses after a 20 sec interval. Cell lysates were then centrifuged at 3000 rpm for 3 min at RT to remove the bulk of chromatin. The supernatant was then incubated with 1.6 μg of anti-DNA-PKcs pS2056 antibodies (abcam 124918) overnight at 4°C, followed by incubation with 25 μl Protein A/G magnetic beads (Pierce) at RT for 2 h. Beads were then washed sequentially once in low salt wash buffer (0.1% SDS, 1% Triton X-100, 2 mM EDTA, 20 mM Tris-HCl pH 8.0, 150 mM NaCl), once in high salt wash buffer (0.1% SDS, 1% Triton X-100, 2 mM EDTA, 20 mM Tris-HCl pH 8.0, 500 mM NaCl), once in LiCl wash buffer (0.25 M LiCl, 1% NP-40, 1% Sodium Deoxycholate, 1 mM EDTA, 10 mM Tris-HCl pH 8.0). Beads were then resuspended in TE buffer (10 mM Tris pH 8.0, 1 mM EDTA) and transferred to a fresh tube and finally eluted with 100 μl elution buffer (1% SDS, 100mM NaHCO_3_). Crosslinks were reverted for the elutions (65°C for 24 h) and DNA was purified with a Qiagen Nucleotide Clean up kit. DNA fragments <300 bp from ChIP elutions were separated by pulling down larger fragments using paramagnetic Ampure XP beads: 65 μl of Ampure XP beads were added to the uncrosslinked ChIP elution and mixed thoroughly. After 10 min incubation at RT, beads were isolated using a magnet. The supernatant was collected and 25 μl of fresh Ampure XP beads were added and mixed thoroughly. After 10 min at RT, beads were separated and the supernatant was cleaned using Qiagen Nucleotide Clean up kit. At the final step DNA was eluted in 60 μl of elution buffer and used for qPCR. 4 different AsiSI sites were monitored using upstream and downstream primers within 40-52 bp from the AsiSI cut site (Supplementary Table 1). %DNA was calculated using equation 2^(Ct(input) – Ct(test))^ x 100%. DNA values obtained for IP in absence of antibodies were subtracted from the values obtained in presence of antibody and used for plotting the graphs in Fig. 5. To monitor the Mre11 nuclease dependence, we treated the human U2OS cells expressing inducible AsiSI with 10 μM NU7441, 100 μM PFM03 and 600 nM 4OHT simultaneously for 1 hour at 37°C before harvesting cells.

## Acknowledgements

We thank Davide Moiani and John Tainer for the Mre11 inhibitors, and members of the Paull laboratory for useful discussions and for critical reading of the manuscript. This work was supported by the National Institute of General Medical Sciences of the National Institutes of Health (GM120554 to I.J.F.), the National Cancer Institute of the National Institutes of Health (CA092584 to I.J.F. and S.P.L.M.), CPRIT (R1214 to I.J.F., RP110465 to T.T.P.), and the Welch Foundation (F-l808 to I.J.F.). I.J.F. is a CPRIT Scholar in Cancer Research. L.R.M. is supported by the National Cancer Institute of the National Institutes of Health (F99CA212452). M.M.S. is supported by a Postdoctoral Fellowship, PF-17-169-01-DMC, from the American Cancer Society. The content is solely the responsibility of the authors and does not necessarily represent the official views of the National Institutes of Health.

## Author contributions

R.A.D., L.R.M., and M.M.S. conceived of and performed experiments and contributed to writing and editing the manuscript; N.M., L.L., and S.P.L-M. provided essential reagents for the experiments and contributed to the editing of the manuscript; I.J.F. and T.T.P contributed to the ideas underlying the experiments and to the writing and editing of the manuscript.

## Data Availibility

All data needed to evaluate the conclusions in the paper are present in the paper and/or the Supplementary Materials. Additional data related to this paper may be requested from the authors.

## List of Supplementary Materials

Figures S1 to S5, and Table S1.

## Supplementary Materials

**Figure S1.**
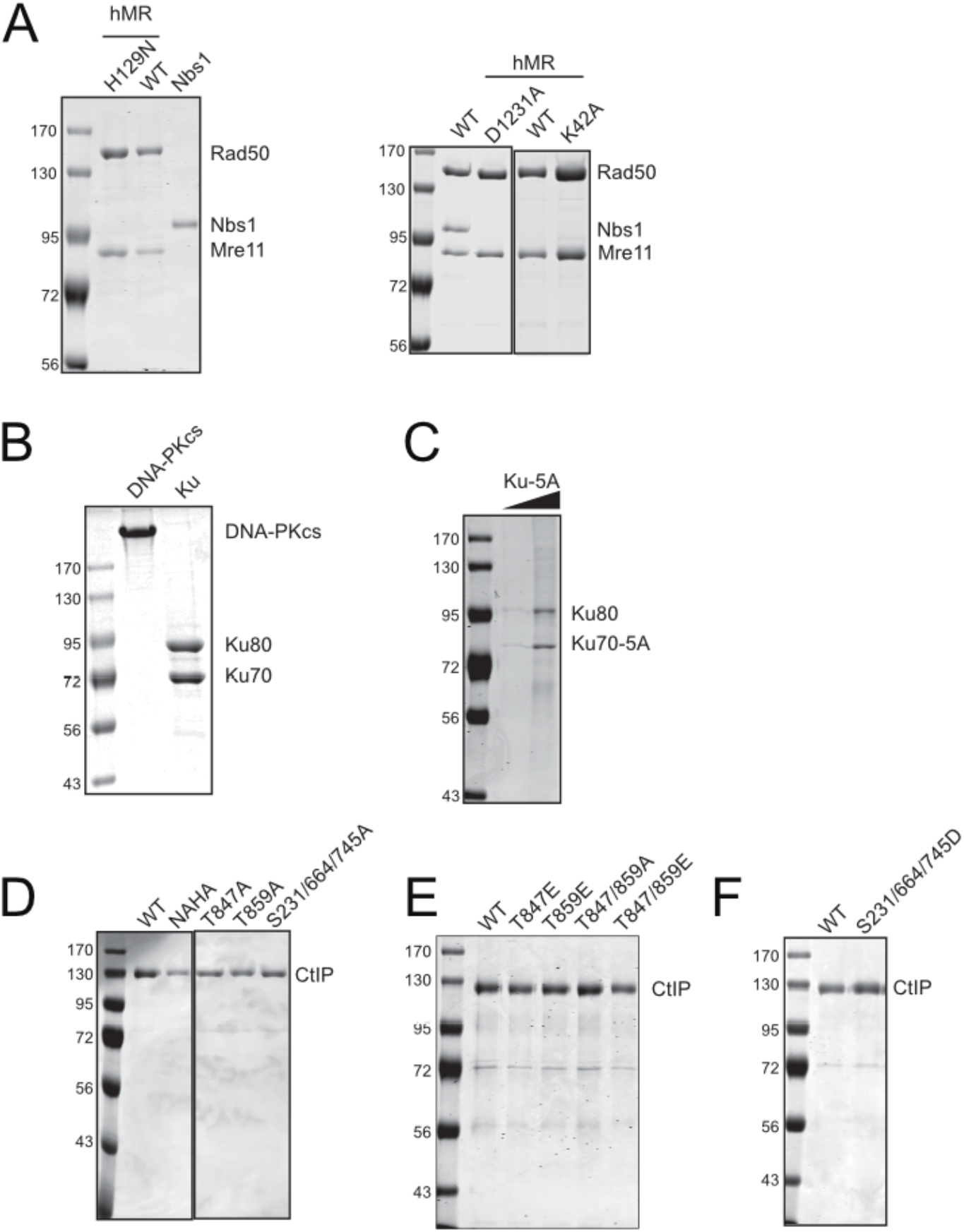
Purified recombinant proteins used in this study. Human recombinant Mre11/Rad50 (hMR) and Nbs1 and hMR(K42A) and hMR(D1231A) mutants (A), Ku70/80 wild-type and DNA-PKcs (B), Ku70(5A)/Ku80 (C), and CtIP proteins (D, E, F) were separated by SDS-PAGE and stained with Coomassie blue. All proteins shown here were purified from insect cells (See materials and methods for details) except DNA-PKcs which is native protein purified from human cells.

**Figure S2.**
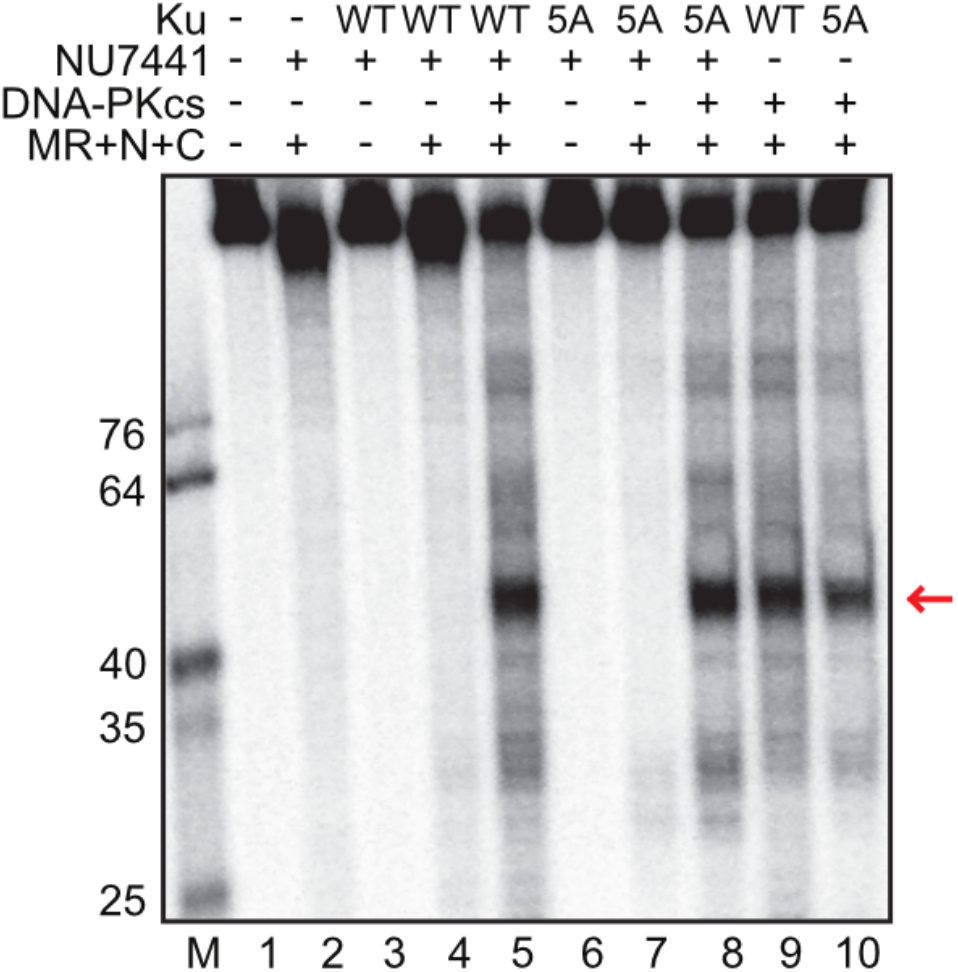
Ku(5A) promotes MRN-dependent endonuclease activity. Nuclease assays were performed as in Fig. 1 in the presence of both magnesium and manganese with wild-type MRN, CtIP, DNA-PKcs, NU7441, and with either wild-type (WT) or 5A mutant (5A) versions of Ku as indicated. Red arrow indicates predominant DNA-PK-dependent MRN product.

**Figure S3.**
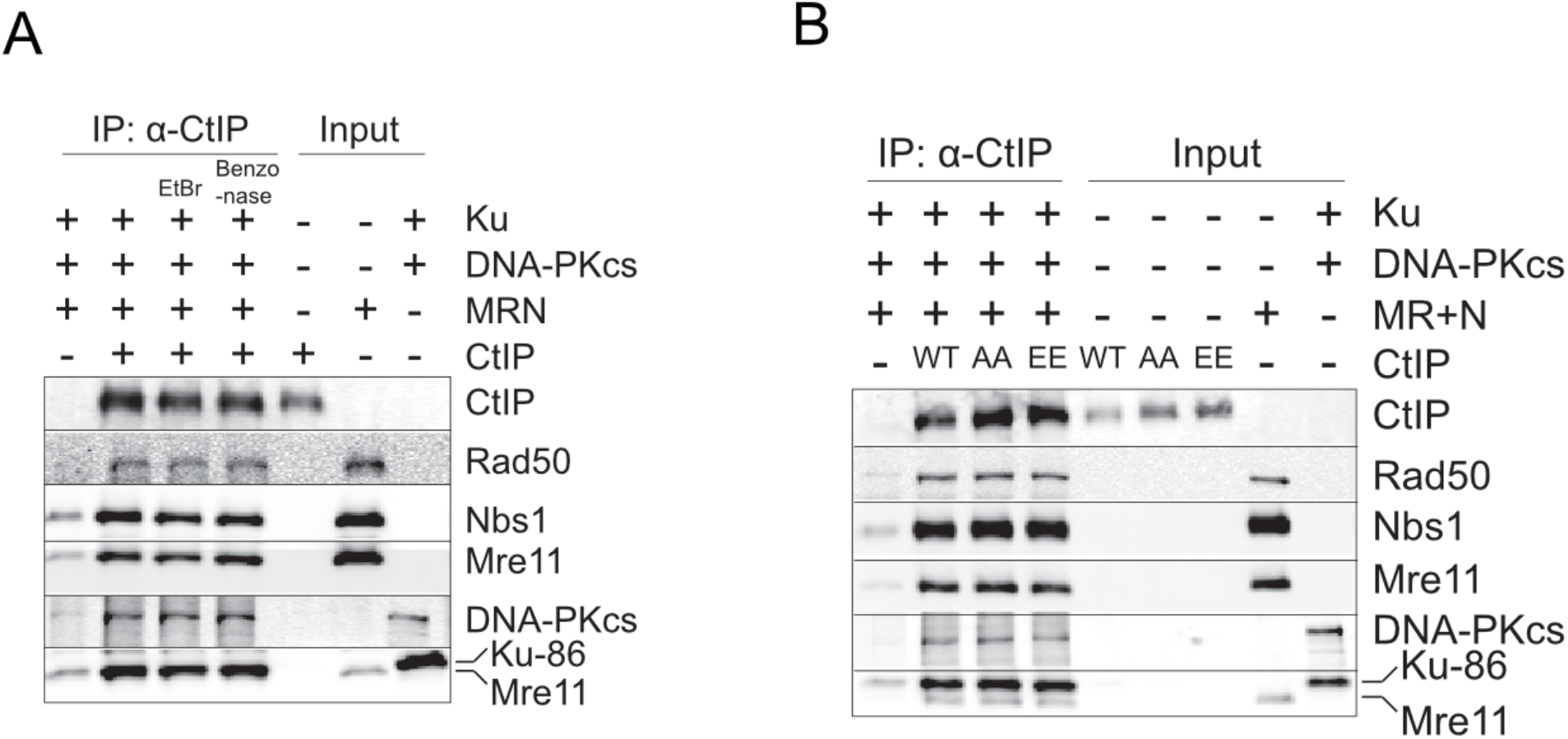
Interactions between CtIP, DNA-PKcs, Ku, and MRN do not depend on DNA or CtIP phosphorylation. (A) Ethidium bromide (5 μg/ml) or Benzonase (1.25 kU) was added to the reactions as indicated. Recombinant proteins were incubated and CtIP isolated by immunoprecipitation as in Fig. 1D. Bound factors were monitored by western blot as indicated. (B) Binding reactions were performed with MRN, DNA-PKcs, Ku, and CtIP wild-type (WT) or mutant T847A/T859A (AA) or T847E/T859E (EE) proteins as in (A).

**Figure S4.**
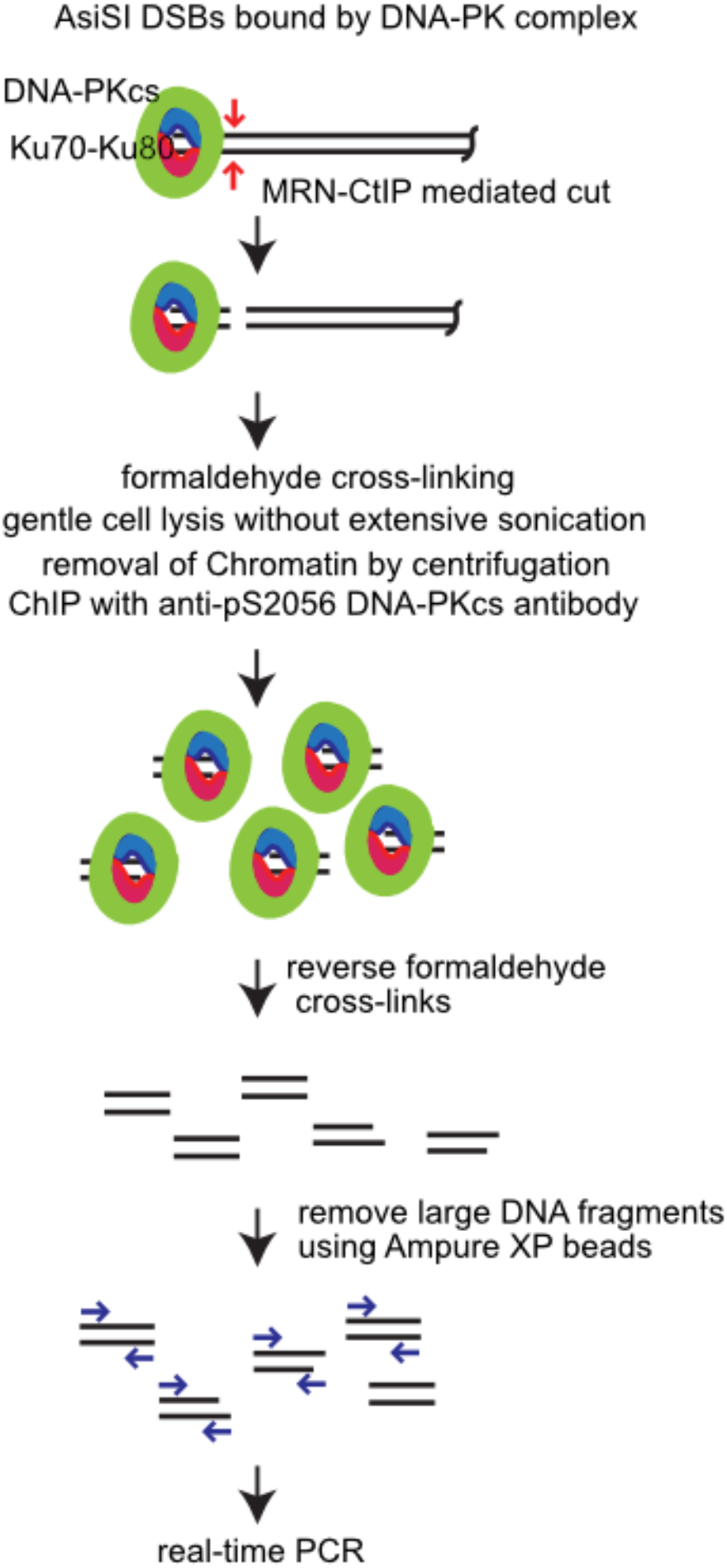
Schematic diagram of GLASS-ChIP protocol. See Materials and Methods for detailed procedure. AsiSI DSBs created in U2OS cells are bound by DNA-PK. Small DNA fragments bound by DNA-PK are generated through endonucleolytic cleavage by MRN with stimulation by CtIP. These protein-bound fragments were stabilized by standard formaldehyde cross-linking but gentle cell lysis was used without excessive sonication in order to avoid fragmentation of genomic DNA. After removal of bulk chromatin, ChIP was performed using anti-DNA-PKcs-pS2056 antibody. After reversal of cross-links and size selection with Ampure XP beads, these DNA fragments were quantified using real-time PCR.

**Figure S5.**
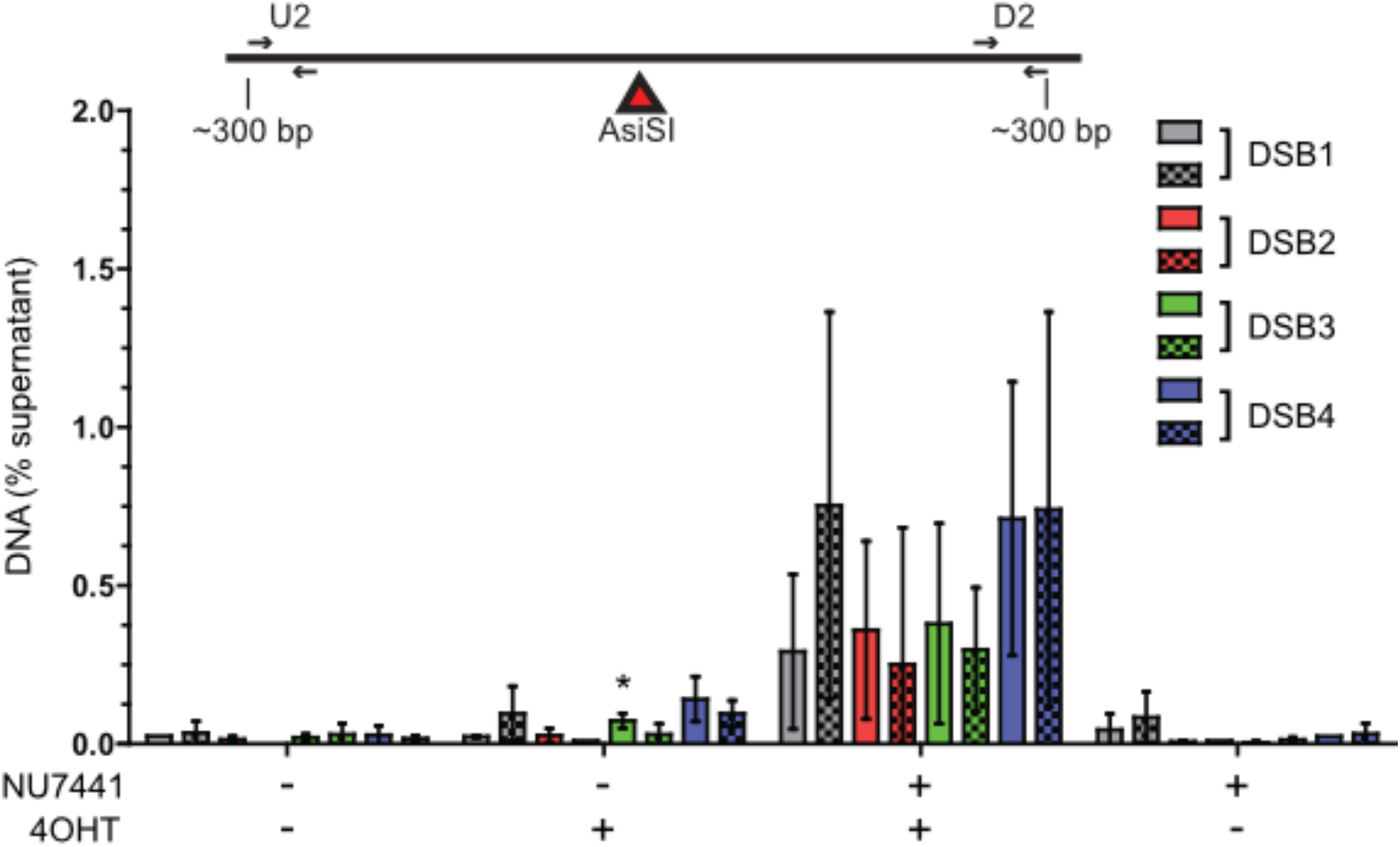
Quantification of DNA immunoprecipitated with anti-DNA-PKcs-pS2056 using primers 300 bp away from AsiSI DSB. Small dsDNA products resulting from nucleolytic cleavage of DNA-PK bound AsiSI-generated DNA ends (dashed line box in Figure 4E) were isolated from U2OS cells using the GLASS-ChIP protocol as described in Fig. S4 and quantified by qPCR using primers located ~300 nt from the AsiSI cut site. Primer set U2 (solid) is upstream whereas D2 (checkered) is downstream of the AsiSI cut sites. The DNA quantitated from U2OS cells in the presence or absence of 4OHT to induce AsiSI and DNA-PKcs inhibitor (NU7441) is shown for 4 AsiSI sites as described in main text. Results shown are from 3 independent biological replicates with student 2-tailed T test performed; * indicates p < 0.05 in comparison to equivalent samples without 4OHT.

**Table S1.**
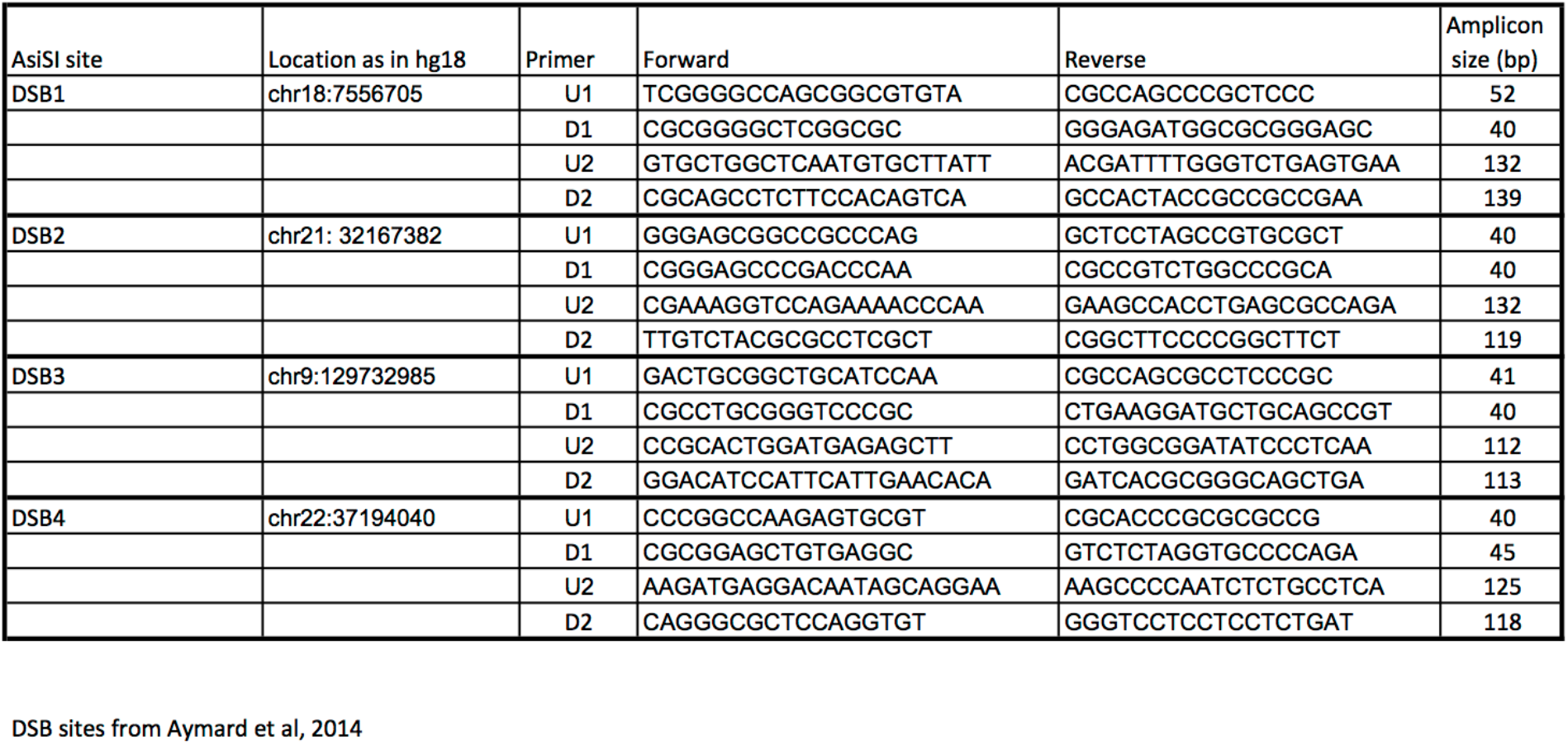
Primers used for qPCR in Fig. 5 and Fig. S5.

